# Radip light-induced phosphorylation changes in microtubule related proteins in arabidopsis

**DOI:** 10.1101/2023.05.30.542878

**Authors:** Denise Soledad Arico, Diego Leonardo Wengier, Natalia B. Burachik, María Agustina Mazzella

## Abstract

Rapid hypocotyl elongation allows buried seedlings to reach the surface, where light triggers de-etiolation and inhibits hypocotyl growth mainly by phytochromes A, B and cryptochromes 1, 2. Dynamic phosphorylation/dephosphorylation events provide a mechanism to rapidly transduce light signals. Only recently we have begun to uncover the earliest phospho-signaling responders to light.

Here, we report a large-scale phosphoproteomic analysis and identify 20 proteins that change their phosphorylation pattern after 20 min of white light pulse compared to darkness. Microtubule-associated proteins (MAPs) were highly overrepresented in this group. Among them, we studied CIP7 (COP1-INTERACTING-PROTEIN-7), which presented microtubule (MT) localization, in contrast to what was previously described. Phosphorylated isoform in Serine 915 (Sp^915^) of CIP7 was detected in etiolated seedlings but undetectable after a light pulse in the presence of photoreceptors, while its expression decays with long light exposure.

The short hypocotyl phenotype and rearrangement of MTs in etiolated *cip7* mutants are complemented by CIP7-YFP and the phospho-mimetic CIP7^S915D^-YFP, but not the phospho-null CIP7^S915A^-YFP suggesting that Sp^915^CIP7 is the active isoform that promotes hypocotyl elongation thorough MT reorganisation in darkness.

Our results reveal that the small repertory of proteins that changes the phosphorylation status after a rapid light signal is tightly focused on MAPs; suggesting that phospho-regulation of microtubule-base processes are early targets during de-etiolation. The evidence on Sp^915^CIP7 supports this idea.

## Introduction

Buried seedlings grow heterotrophically following the skotomorphogenic program of development, a pattern that includes fast hypocotyl elongation in dark-grown seedlings of Arabidopsis. PHYTOCHROME-INTERACTING FACTORS (PIFs) transcription factors promote the expression of genes involved in the elongation of hypocotyl cells (Leivar & Monte, 2014). In addition, the E3 ubiquitin ligase CONSTITUTIVE PHOTOMORPHOGENIC 1 (COP1) promotes proteome-mediated degradation of transcription factors such as ELONGATED HYPOCOTYL 5 (HY5) (Osterlund *et al*., 2000) which, represses hypocotyl cell elongation (Gangappa & Botto, 2016). When shoot organs emerge from the soil, light is perceived mainly by the red far-red light phytochromes A and B (phyA and phyB) and the blue UV-A light cryptochromes 1 and 2 (cry1 and cry2) (Kami *et al*., 2010), the four most important photoreceptors involved in the shift from skoto-to photo-morphogenesis de-etiolation (Mazzella & Casal, 2001; Fox *et al*., 2015). Light-activated photoreceptors repress COP1 and PIFs activities (Martínez *et al*., 2018) arresting hypocotyl cell elongation (Sibout *et al*., 2006).

Cell expansion depends on turgor pressure and cell walls loosening (Feng *et al*., 2016; Cosgrove, 2018). Plant cell architecture and expansion are controlled by events that occur at the interface of the plasma membrane and the cell wall, including vesicle trafficking and the control of microtubule (MTs)-mediated cell wall deposition (Endler *et al*., 2015; Feng *et al*., 2016). Cortical MTs regulate cell expansion by setting the orientation of cellulose microfibrils in the cell wall (Lloyd, 2011). In the dark, where hypocotyl cell elongation is maximal, the cortical MTs that form the intracellular cytoskeleton are oriented perpendicular (or transverse) to the growth axis directing the deposition of cellulose microfibrils that favours cell elongation (Elliott & Shaw, 2018). Upon exposure to light, MTs rearrange obliquely or longitudinally in the hypocotyl cells and growth is inhibited (Li *et al*., 2011; Sambade *et al*., 2012) in approximately 15 min (Lindeboom *et al*., 2019). However, the links between light and MT dynamics remain elusive.

Light induces changes in protein phosphorylation (Inoue *et al*., 2008; Deng *et al*., 2014; Shin *et al*., 2016; Liu *et al*., 2017; Wang *et al*., 2021; Gao *et al*., 2022). Some of them are relevant to the control of hypocotyl growth such as phytochrome-mediated phosphorylation of PIFs, which triggers their degradation (Shen *et al*., 2008; Ni *et al*., 2014, 2017). However, our knowledge of light-induced changes in proteome-wide phosphorylation are limited. The analysis of the phosphoproteome in etiolated maize seedlings exposed to light has revealed important modifications in proteins involved in photosynthesis (Gao *et al*., 2020). However, the focus of that study is on medium-long term changes (1-12 h of light exposure). Thus, we ignore the earliest cellular functions targeted by phosphorylation events, which can certainly take place within a few minutes.

Here, we present a large-scale phosphoproteome study of dark-grown Arabidopsis seedlings of the wild type and of the *phyA phyB cry1 cry2* quadruple mutant exposed to 20 min of white light (WL). We show that microtubule associated proteins (MAPs) are strongly overrepresented among the proteins showing rapid changes in phosphorylation. Among them we identify COP1-INTERACTING PROTEIN 7 (CIP7) (Yamamoto *et al*., 1998) as a MAP, which phosphorylated isoform promotes hypocotyl elongation and favours transverse MTs orientation in etiolated seedlings.

## Materials and Methods

### Phosphoproteomic assay

#### Plant Material

*Arabidopsis thaliana* WT seeds from ecotype *Landsberg erecta* were obtained from the Arabidopsis Biological Resource center (ABRC, Ohio, USA). No specific permission is needed to use WT ABRC seeds. The quadruple mutant *phyA-201 phyB-1 cry1-1 fha-1* named *phyAphyBcry1cry2* was generated by Mazzella & Casal, 2001. Seeds were sterilized for 2 h in a Cl_2_ (g) atmosphere generated by the addition of 1.5 ml HCl (37% v/v) to 50 ml of commercial bleach (8.25% sodium hypochlorite). Sterilized seeds were sown on 0.5X MS (0.5X Murashige-Skoog basal salts buffered with 0.5g/L MES pH5.8 and 0.8% (w/v) agar) in petri dishes and incubated at 4 °C in darkness during 3 d for stratification. Chilled seeds were exposed to a 2 h red light pulse (100 μmol m^−2^s^−1^) at 22°C to synchronize germination, and then kept in darkness for 5 d before harvest. 3 conditions were analyzed (Supplemental figure 1): **WTD**: 5-d-dark-grown WT seedlings. **WTL**: 5-d-dark-grown WT seedlings exposed to a 20-min WL pulse (100 μmol m^−2^s^−1^) before harvest. ***phyAphyBcry1cry2*L**: 5-d-dark-grown quadruple mutant seedlings exposed to a 20-min WL pulse (100 μmol m^−2^s^−1^) before harvest.

#### Procedure and data acquisition

Total protein extraction, trypsin digestion of proteins, phosphopeptides enrichment and LC-MS/MS (Supplemental figure 1) is fully described in Arico *et al*., 2021. Three biological replicates per condition were analyzed. The identification of phosphopeptides was performed using the Max-Quant/Andromeda engine, setting as fixed modifications: Carbamidomethylation (C), and variable modifications: Oxidation (M) + Phosphorylation (STY). The database employed was UniProt CP_Arabidopsis Thaliana. 1% FDR filter was applied. The quantification was done using signal intensity values of peptides. A peptide was considered quantifiable if it had at least 2 signal intensity values among the three biological replicates. Missing signal intensity values were imputed with a noise value corresponding to the 1-percentile of all intensity values by sample. The mass spectrometry proteomics data have been deposited to the ProteomeXchange Consortium via the PRIDE (Vizcaíno *et al*., 2016) partner repository with the dataset identifier PXD008274.

#### Differential phosphorylation analysis

In order to identify the light-induced phosphopeptides, the abundance of each phosphopeptide was compared between WTL and WTD. The abundance was computed as an average of 3 independent signal intensity values per phosphopeptide quantified in each condition. A phosphopeptide was considered significantly enriched in WTL vs WTD if it simultaneously fulfilled a Welch’s p-value < 0.05 and a Z-score absolute value > 1.96. In order to identify the photoreceptor dependency in response to light, the abundance of each phosphopeptide was compared between *phyAphyBcry1cry2*L and WTL. Equally, the difference was significant if the Welch’s p-value < 0.05 and a Z-score absolute value > 1.96.

### Generation of constructs and transgenic lines

The Gateway cloning strategy was used to make the transcriptional reporters, translational reporters, truncated versions, CRISPR-Cas9 mutants and phosphomimetic/null isoforms of *CIP7*; according to the protocols in the Gateway Cloning Technology booklet (Invitrogen, Carlsbad, CA). Amplifications were performed with Q5 enzyme (New England Biolabs, Massachusetts, USA.) using genomic DNA of *Arabidopsis thaliana* as a template. The details of the primers are specified in Table S1.

The PCR products and the pENTR-D-TOPO vector were digested using the restriction enzymes NotI-HF y AscI (New England BioLabs) following the instructions of the supplier and the digested vector was dephosphorylated by TSAP enzyme to avoid its recircularization (Promega, Madison, WI, USA). After electrophoresis, all the fragments were purified using QIAEXII Gel Extraction Kit (QIAGEN, Germany). Ligation was performed using T4 ligase enzyme (Promega, Madison, WI, USA) and a molarity ratio of insert:vector equal to 3:1 for all the constructs. 20-100 ng of vector and 10U of enzyme were used in a 10μl final reaction volume. The reaction took place overnight at 16°C.

Once the entry vectors carrying the constructs were confirmed by sequencing, the binary vectors were generated by LR recombination between pENTR-D-TOPO and the destiny vectors detailed in Table S2; according to the protocols in the Gateway Cloning Technology booklet (Invitrogen, California, USA). All constructs were sequenced by Macrogen Inc., South Korea; and analyzed using blast.ncbi.nlm.nih.gov/Blast.cgi.

Arabidopsis was transformed following floral dipping and T1s seeds were selected on the appropriate antibiotic according to the Arabidopsis Handbook standard protocols. Transformants in WT ecotype Col-0 and mutant *cip7_ed2* backgrounds, were selected in half-strength MS-agar 0.8% dishes supplemented with the antibiotic specified in Table S2. The plants transformed with the microtubule marker *UBQ10::MBD-mCherry* (Ivanov & Harrison, 2014) were selected also in half-strength MS-agar 0.8% dishes supplemented with Kanamycin 50mg/l. 2 independent homozygous lines were generated for each construct per plant genetic background.

#### Generation of CRISPR-Cas9 mutants

The strategy used was based on the protocol from Miao *et al*., 2013. The target portion of RNA guides for leading CAS9 endonuclease to the 5’-end of *CIP7*, were designed using CRISPR-P2.0 (crispr.hzau.edu.cn/CRISPR2/). Forward oligonucleotides contained 4 compatible bases to the overhangs of BsaI; an additional G that has been added after the overhang so that any target sequence from CRISPR-P2.0 can be used (both those that start with G or A); and the 20-base target (starting with G or A). Reverse oligonucleotides contained 4 compatible bases to the overhangs of BsaI; the 20-base target; and an additional C at the end due to the extra G in the forward oligonucleotides. The sequences can be found in Table S1.

Guide RNAs were cloned by ligation of a double stranded 20 DNA fragment harbouring 4-base overhangs on each end compatible to overhangs on a pENTR vector carrying Arabidopsis *U6* promoter and guide RNA scaffold, and a reporter cassette that expresses *PIP2A:mCherry* under the control of *35S*. Confirmed constructs were recombined to M3T-UBQp-CAS9-GW. Arabidopsis WT plants were transformed and selection was carried out in half-strength MS-agar 0.8% dishes supplemented with Kanamycin 50mg/l. T1s were genotyped using primers cCIP7F and CRISPR CR6 rv (Table S1). The editions were corroborated by Sanger sequencing. T2s that lacked the *mCherry* cassette (and thus the CRISPR/Cas9 constructs) were selected for further analysis. 2 independent homozygous mutant lines were generated and corroborated by sequencing.

#### Generation of phospho-mimetic and phospho-null isoforms of CIP7

The phospho-mimic and phospho-null versions of CIP7 were obtained by site-directed mutagenesis using primer pairs detailed in Supplemental Table 1. The pENTR-D-TOPO carrying CIP7 gene (without stop codon) was used as template for substituting Sp^915^ to D and A respectively. The degradation of the template carrying the WT version of CIP7, was performed with DpnI enzyme, following the instructions of the manufacturer (New England Biolabs Inc., Massachusetts, USA). Transformation of *E. coli*, LR recombination with the destiny vector pH35GY, transformation of *Agrobacterium tumefaciens*, and transformation of plants were performed as previously described.

### GUS histological assay

GUS histological assays were performed according to Jefferson *et al*., 1987 using dark-grown *pCIP7::GUS* seedlings exposed to light treatments specified in each experiment. A promoter of 2500pb was used. *pBG GUS* seedlings, carrying the empty vector, were used as controls. Staining proceeded for 24 hours at 37°C in the dark. Seedlings were washed twice with ethanol 70% and twice with water. Samples were mounted in Hoyer’s medium (Anderson, 1954) for imaging in a Leica DM2500 microscope using Differential Interference Contrast (DIC).

### RT-qPCR

200 mg of seedlings per sample were grinded and freezed with N_2_ (l). Total RNA extraction was performed using RNeasy Plant Mini Kit (QIAGEN, Germany). RNA integrity was corroborated by gel electrophoresis and quantified by Nanodrop. 2μg of RNA per sample were treated with DNase RQ1 RNase-Free (Promega, Madison, WI, USA); and used for synthesis of cDNA with Super-Script IV enzyme (Invitrogen), in a 20 μl reaction using oligo (dT) primers. cDNA samples were diluted 1:10 and 5 μl were taken for quantitative real time PCR (qPCR). qPCR reactions were performed in a DNA Engine Opticon 2 System (MJ Research, USA) using the 5x HOT FIREPol EvaGreen® qPCR Mix Plus (NO ROX) kit (Solis BioDyne, Estonia). Primers RT-qPCR2 cip7 F and RT-qPCR2 cip7 R (Table S1) were designed with primer3plus.com. EF-1α (Elongation Factor 1α, At5g60390) was used as normalization control (Table S1). The conditions for PCR were optimized with respect to primer concentrations, primer annealing temperatures, and duration of steps. Cycling conditions were 95°C for 15 min followed by 38 cycles of 15 s at 94°C, 12 s at 60°C, and 12 s at 72°C. PCR for each gene fragment was performed alongside standard dilution curves of cDNA pool. All gene fragments were amplified in duplicate from the same RNA preparation, and the mean value was considered for each replicate. Data are means of 2 independent experiments.

### Confocal microscopy assays

#### Image acquisition

CIP7 translational reporters seedlings were imaged in a Zeiss LSM 880 Indimo AxioObserver inverted microscope using a dry objective 20x/NA 0.8 (or 40x/NA 1.2 for MTs orientation measurements); with the following settings to detect YFP signal: Excitation laser, 514nm; emission λ, 542nm; gain, 680. The image size was 1024 x 1024 pixels and the pixel size: 0.2μm x 0.2μm. The microtubule marker *UBQ10::MBD-mCherry* was imaged with the following setting: Excitation laser, 543nm; emission λ, 653nm; gain, 950. Single images were acquired in all cases, except for Supplemental Figure 3 in which z-stacks of 32.4 μm in 18 steps were taken starting from the most external periclinal wall of epidermal cells. For light grown seedlings, an additional channel was configured for detecting the autofluorescence of chlorophyll in the λ emission range of 600nm-700nm. The top half/upper-hypocotyl of at least 3 plants of each genotype were imaged per experiment and each experiment was repeated at least 4 times. All the seedlings were mounted in water for image acquisition; except for the Oryzalin assay in which 3 d etiolated seedlings were incubated for 30 minutes in 10μM Oryzalin (stock 10mM in ethanol) or Ethanol mock and mounted in the respective solution. For colocalization assays, CIP7 translational reporters were crossed with the microtubule marker and F1 generation was imaged as previously described for both fluorescent proteins (YFP and mCherry, respectively). For Figure 2B and Supplemental Figure 4B, images were taken with the Airyscan Super-resolution detector in an unidirectional scan, using an immersive objective 40x/NA= 1.2 (gain= 950, pinhole 1.55 UA). The images size was 992 x 992 pixels and the pixel size: 0.05 μm x 0.05μm. The quantitative colocalization was done in ImageJ/Fiji (Schindelin *et al*., 2012) using JACoP (Just Another Colocalization Plugin) applying manual threshold. Pearson’s and Mander’s coefficients were obtained.

#### Quantification of images

Estimation of CIP7-YFP levels by fluorescence intensity quantification was done with ImageJ/Fiji. In all cases, single images of top half/upper-hypocotyls were acquired with the setting for YFP described above. Measurements were performed on cortical microtubules located towards the external periclinal cell wall of epidermal cells. A ROI of 10 μm x 20μm was selected to measure mean grey values. 3 to 6 randomly selected cells were measured per seedling and 3 measurements were done per cell. At least 200 cells were measured per genotype. Box plots were done using R (R CoreTeam, 2022).

Average orientation angle and anisotropy of cortical microtubules were measured using the FibrilTool plug-in designed for ImageJ/Fiji (Boudaoud *et al*., 2014). In order to compare the different genotypes, all the translational reporters and controls were crossed to *UBQ10::MBD-mCherry*, and imaged using the acquisition setting previously described for mCherry. Measurements were performed on cortical microtubules located towards the most external periclinal cell wall of hypocotyls’ epidermal cells. 3 to 6 cells were measured per seedling, selecting ROIs with the size of the cell. At least 100 cells were measured per genotype in each experiment. Experiments were repeated at least 8 times. Average angles with anisotropy values higher than 0.05 were organized in three categories: transverse (0°-30°), oblique (30°-60°) and longitudinal (60°-90°) (Fig. S8).Those with anisotropy values minor to 0.05 were considered to be random. For each genotype, absolute frequency was calculated per category and illustrated with a bar plot using Graphpad Software. Significant differences among frequencies were analyzed using a one-way or two-way analysis of variance and Tuckey test.

### In-silico analysis

Identification of potential microtubule associated proteins (MAP) among the light-induced differential phosphoproteins was carried out with MAPanalyzer (Zhou *et al*., 2015). The list of AGI codes was submitted in systbio.cau.edu.cn/mappred/index.php, and a prediction of microtubule binding motifs in the primary sequence was computed with a specificity higher than 95%. The proteins bearing microtubules binding motifs with “high” and “very high” specificity (< 95% and 99%, respectively) were considered as MAPs. The prediction of microtubules binding motifs present in the primary sequence of CIP7, was done in the same way.

The prediction of the most accurate tridimensional structure of CIP7 was performed with I-TASSER (Zhang, 2008; Roy *et al*., 2010; Yang *et al*., 2015). The amino acid sequence was submitted in https://zhanggroup.org/I-TASSER/.

### Hypocotyl length measurements

Seeds were sown on 0.5X MS on square petri dishes (12cm x 12cm). After stratification and germination induction, they were kept vertically in darkness during the number of days specified in each experiment. Twelve to fifteen seedlings of a maximum of 8 genotypes were sown in each plate at a time. The plates were imaged and the hypocotyl length was measured using the segmented line tool of ImageJ/Fiji. An average of 10 seedlings’ measurements per genotype was considered as a biological replicate. Between 3 and 6 biological replicates per genotype were analyzed in each experiment.

Data was analyzed by one way ANOVA followed by *a posteriori* Tuckey test for multiple comparisons. Homoscedasticity was evaluated by Levene’s test and normality by modified Shapiro-Wilks test. Plots depicted mean ± standard error and statistics were done using GraphPad Prism versión 6.00 (GraphPad Software, La Jolla California, USA).

## Results

### MAPs are overrepresented in the early light-induced phosphoproteome

To identify early light-responsive proteins that change their phosphorylation status, and discern if there is dependency on the four main photoreceptors (phyA, phyB, cry1 and/or cry2); we conducted a large-scale mass spectrometry-based (LC-MS/MS) quantitative phosphoproteome assay in Arabidopsis. We analyzed samples obtained from WT seedlings grown during 5 d in complete darkness (WTD) or that received a 20 min of white-light pulse before harvest (WTL); and *phyA phyB cry1 cry2* quadruple mutants grown for 5 d in darkness and exposed to a 20 min pulse of white-light before harvest (*phyAphyBcry1cry2*L) (Fig. S1).

In total, 2160 phosphopeptides were identified representing 1098 unique phosphoproteins at a 1% FDR. The number of phosphoproteins obtained in our work is comparable with those obtained using similar strategies (Waterworth *et al*., 2019; Kamal *et al*., 2020). To identify the proteins that changed their phosphorylation status upon light, we compared the abundance of each phosphopeptide between WTD and WTL treatments. LogWTL/WTD = 0 indicates that no changes in the phosphorylation status were detected; logWTL/WTD > 0 indicates phosphorylation status in response to light, while logWTL/WTD < 0 suggests dephosphorylation status in response to light. We computed a Welch statistical test with a significant cut-off value of p < 0.05 and a z-score absolute value < 1.96 to discard false positives (see M&M) (Table 1). The volcano plot combines the statistical significance with the fold-change (Fig. 1). After 20 min of WL treatments, twenty-two phosphopeptides corresponding to 20 unique phosphoproteins changed their phosphorylation status significantly. This resulted in 17 phosphorylated proteins and 3 non-phosphorylated proteins in response to light (Fig. 1 and Table 1). Many of the phosphosites identified in this study have already been reported as early light-induced ones: CRY2 is phosphorylated by light in Yp^604^ (p-value = 0.017) (Table 1) (Liu *et al*., 2017), while PHOT1 is phosphorylated in Sp^13^ (p-value = 0.014), Sp^23^ (p-valor = 0.008), Sp^382^ (p-valor = 0.051) and Sp^409^ (p-valor = 0.022) (Table 1) (Inoue *et al*., 2008; Sullivan *et al*., 2008; Deng *et al*., 2014).

**Fig. 1.**
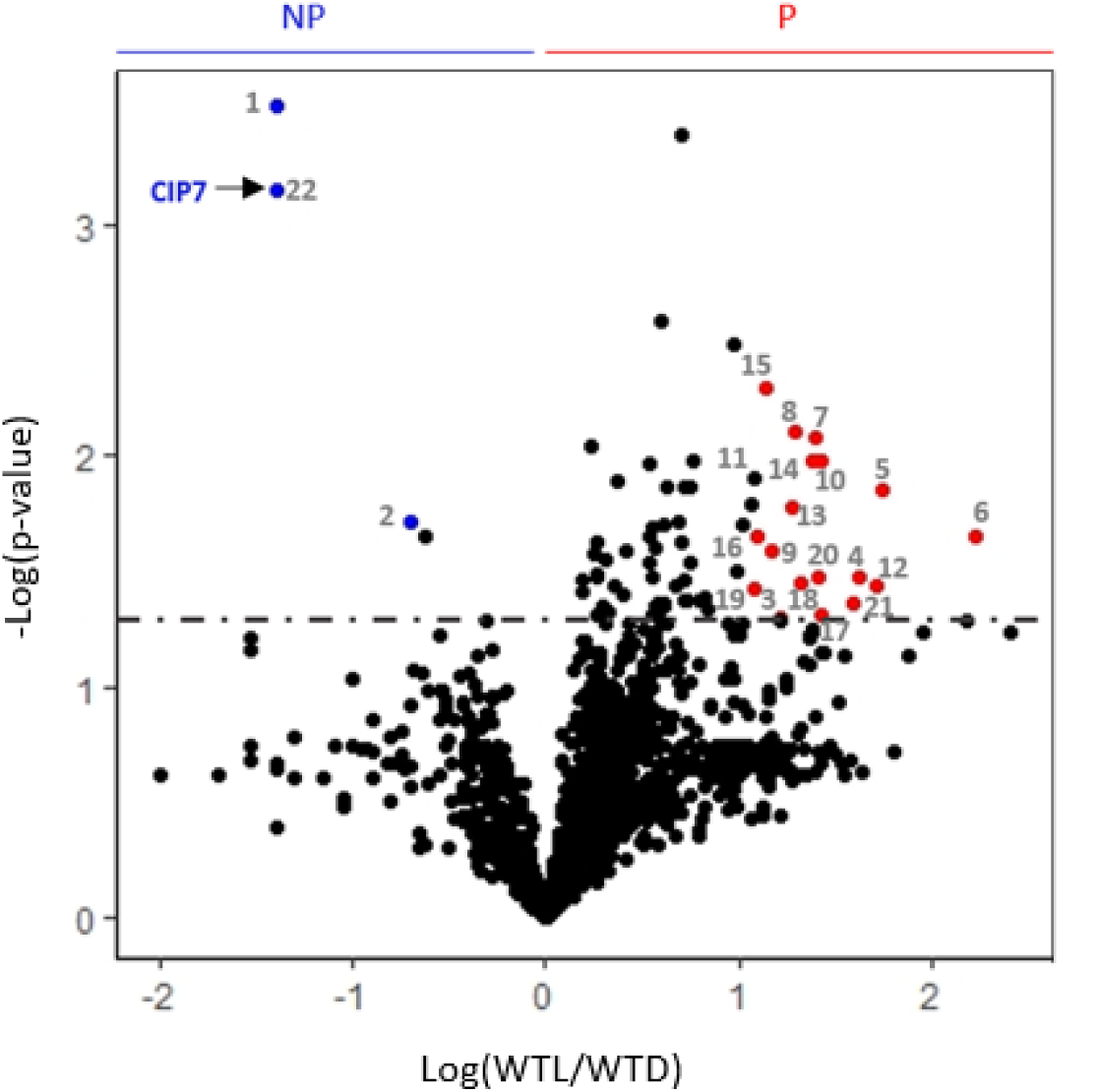
Early light-induced changes in phosphopeptide phosphorylation. Volcano plot depicting WTL/WTD ratios and the corresponding Welch’s test p-value in logarithmic scale for each phosphopeptide identified. Threshold of significance (dashed line): p value < 0.05 and z-score > |1.96|. Twenty two phosphopeptides change their phosphorylation pattern upon 20 min of WL exposure. NV: Non-Variable phosphopeptides, in black; NP: Non phosphorylated phosphopeptides in light, in blue; P: Phosphorylated phosphopeptides in light, in red. Significant phosphopeptides are numbered according to Table 1. CIP7 phosphopeptide is indicated.

**Fig. 2.**
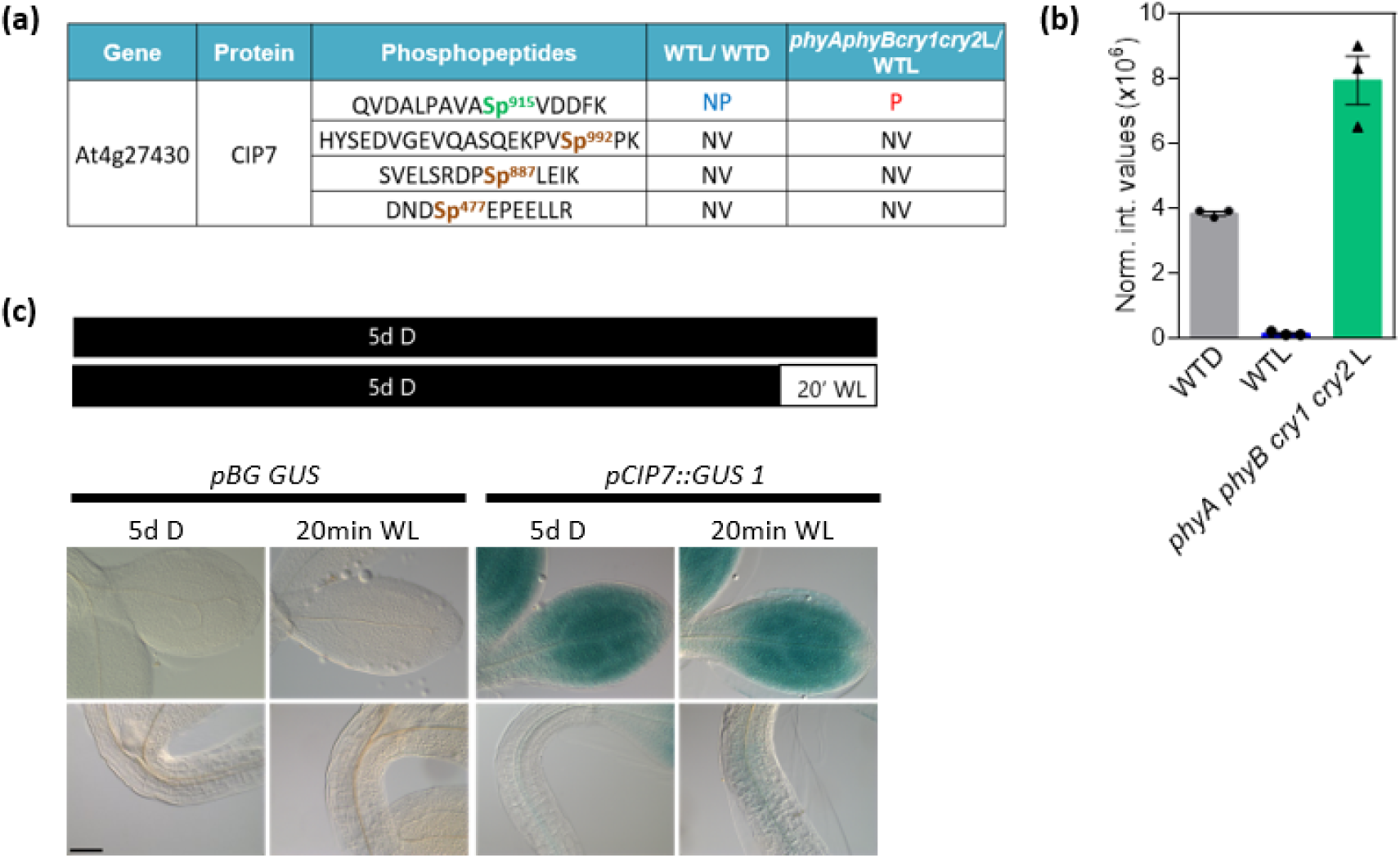
Only the isoform **Sp^915^ of CIP7** is abundant in darkness but undetectable after a light pulse in a photoreceptor dependent manner. **(a)** Phosphopeptides corresponding to CIP7 present in the phosphoproteome. NP: Non-phosphorylated phosphopeptide in WTL, in blue P: Phosphorylated phosphopeptide in *phyA phyB cry1 cry2* L, in red; NV non-variable phosphopeptides, in black. Sp: Phosphorylated Serine. **(b)** Plot depicting the normalized intensity values obtained in the MS/MS spectra analysis for QVDALPAVASp^915^VDDFK in WTD, WTL and *phyA phyB cry1 cry2L*. Mean + SE of 3 independent replicates for each condition. Data was analysed by one-way ANOVA: WTD/WTL Welch’s p-value < 0.01. *phyA phyB cry1 cry2L*/WTL Welch’s p-value < 0.001. **(c)** GUS expression pattern of *pCIP7::GUS* compared to the no promoter control (pBG GUS) in 5-d-old dark-grown seedlings either exposed or not to a 20 min WL pulse (the conditions studied in the phosphoproteomic assay). Scale Bar: 100 μm.

**Table 1.**
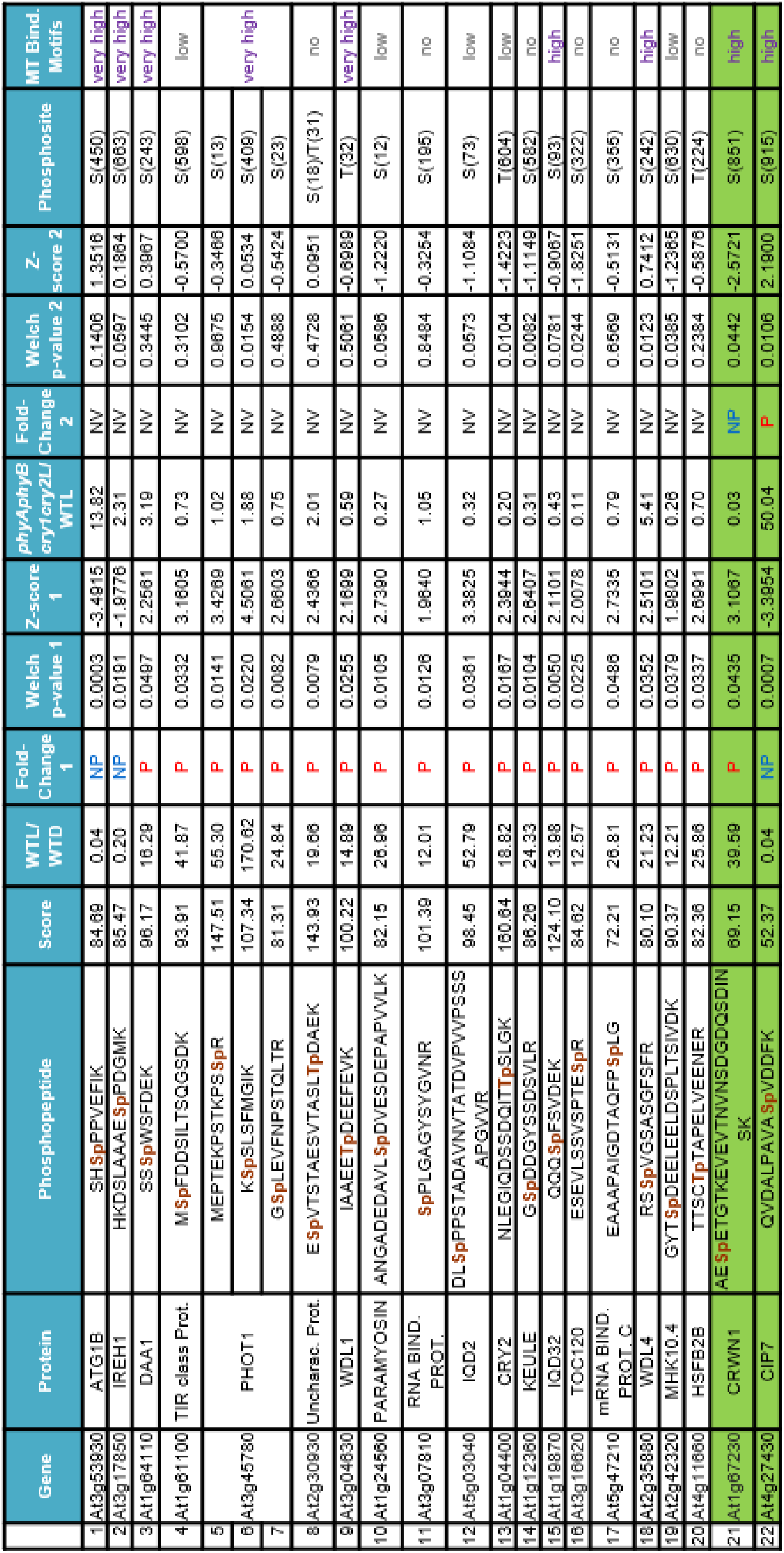
List of light-induced differentially phosphorylated peptides. The abundance of each phosphopeptide was compared between WTL and WTD (Fold Change 1), and *phyAphyBcry1cry2*L and WTL (Fold Change 2). A phosphopeptide was considered as a significant variant if it simultaneously fulfils a Welch’s test p-value < 0.05 and a Z-Score absolute value > 1.96. NV: Non-Variable phosphopeptides, in black; NP: Non-phosphorylated phosphopeptides in light, in blue; P: Phosphorylated phosphopeptides in light, in red. Three independent replicates were analysed for each condition WTD, WTL and *phyAphyBcry1cry2*L. Green box, photoreceptor responsive phosphopeptides. Sp: Phosphorylated Serine. Tp: Phosphorylated Threonine. Phosphosite column indicates the position of the phosphorylated amino acid within the protein. In violet, Microtubule binding motifs prediction with >99% very high, > 95% high or < 95% low specificity according to *MAPanalyzer*.

In a previous study we found that the GO (Gene Ontology) terms “regulation of mRNA splicing” and “cytoplasmic MTs compounds” were specifically enriched in the phosphoproteome of dark-grown Arabidopsis seedlings (Arico *et al*., 2021). None of the 40 unique phosphoproteins related to “regulation of mRNA splicing” identified in darkness (Arico *et al*., 2021) changes their phosphorylation status after 20 min of light. The bioinformatic tool *MAPanalyzer* allows the analysis of known MAPs (Microtubule Associated Proteins) or predicted ones (Zhou *et al*., 2015). According to MAP*analyzer* database, 5% of the genes in the Arabidopsis genome are predicted to have MT binding motifs with a very high (>99%) or high (>95%) specificity threshold (Fig. S2). The frequency of predicted MAPs increased significantly (14%) in the etiolated WT phosphoproteome (Fig. S2). Additionally, this frequency even significantly increased (45%) in the pool of phosphoproteins that changed their phosphorylation pattern after 20 min of light compared to darkness (Fig. S2, Table 1). Association to MTs of WDL1, WDL4 and IQD32 is supported experimentally (Yuen *et al*., 2003; Hamada *et al*., 2013; Liu *et al*., 2013). Among the predicted MAPs in Table 1, PHOT1 is associated with MTs in Arabidopsis in vivo (Kaiserli *et al*., 2009; Sullivan *et al*., 2010), and has recently been described as a cytoskeleton associated protein involved in organelle positioning under light stress (Kumar *et al*., 2023). IREH1 controls root skewing and MTs network (Yue *et al*., 2019); PROTEIN CROWDED NUCLEI 1 (CRWN1) links the nucleoskeleton and cytoskeleton (Sakamoto & Takagi, 2013; Wang *et al*., 2013); and DAA1 belongs to the same meiotic clade of AAA+ proteins such as katanin and spastin which are involved in MT severing and disassembly (Baas *et al*., 2005). No data in the literature associated CIP7 to the cytoskeleton structures.

Our results show that a reduced pool of proteins changes their phosphorylation pattern after a rapid light signal and that significant changes in the phosphorylation status of MAPs occurs after a short light treatment.

### Phosphorylated levels of the isoform Sp^915^ of CIP7 are abundant in darkness and decreased by light-activated sensory receptors

To study if the light-induced changes in the phosphorylation status are dependent or independent of the presence of phyA, phyB, cry1 and/or cry2, we compared the abundance of the 22 light-responsive phosphopeptides between WTL and *phyAphyBcry1cry2*L samples. Twenty of the 22 phosphopeptides did not show statistically different changes in the relation *phyAphyBcry1cry2*L/ WTL, indicating that early light changes in phosphorylation were independent of photoreceptors (Table 1). Only two phosphopeptides showed significant changes, CRWN1 with a relation of *phyAphyBcry1cry2*L/ WTL < 1, and CIP7 with a relation of *phyAphyBcry1cry2*L/ WTL >1 (Table 1). This result suggests that light-induced changes in phosphorylation patterns of CRWN1 and CIP7 depend on the presence of phyA phyB cry1 and/or cry2.

From the proteins that changed their phosphorylation status by light, CIP7 caught our attention for many reasons: First, it has been described as a transcription factor with nuclear localization (Yamamoto *et al*., 1998) but predicted as a MAP with high confidence in our study (Table 1). Second, it physically interacts with the photomorphogenic repressor COP1 (Yamamoto *et al*., 1998). Third, from the four phosphopeptides of CIP7 identified in darkness only the one that contains the Serine 915 (Sp^915^) (QVDALPAVA**Sp^915^**VDDFK) was undetectable after 20 min of light treatment, but present in the *phyA phyB cry1 cry2* quadruple mutants exposed to light (Table 1 and Fig. 2 a,b). This reduction cannot be explained by changes in transcriptional expression, as GUS activity in transgenic plants expressing the promoter of CIP7 fused to the β-glucuronidase (GUS) reporter gene, was readily detected in 5-d-old dark-grown seedlings as well as after 20 min of a WL pulse, in cotyledons and apical regions of the hypocotyl (Fig. 2c). These results suggest that Sp^915^CIP7 isoform is reduced after 20 min of light in a phyA, phyB, cry1 and/or cry2 dependent manner.

### CIP7 localizes to microtubules

We then predicted the presence of MTs motifs in CIP7 using the bioinformatic tool *MAPanalyzer*. CIP7 is a large protein of 1058 amino acids with 24 predicted MT-binding motifs (Fig. S3a,b). Two structural domains have been defined for CIP7: the N-terminal domain that includes the amino acids 1 to 386, and the C-terminal domain that includes the amino acids 387 to 1058 (Yamamoto *et al*., 1998). These domains correlate with the tridimensional structure of the protein predicted with the highest confidence by the protein fold algorithm I-TASSER (Zhang, 2008; Roy *et al*., 2010; Yang *et al*., 2015) (Fig. S3c). We investigated the subcellular localization of CIP7 in transgenic lines expressing the *p35S::CIP7-YFP*, *p35S::YFP-CIP7* and the *pCIP7::CIP7-YFP* constructions. In 3-d-old dark-grown seedlings, *p35S::YFP* transgenic control lines displayed detectable fluorescence in nuclei and cytoplasm of hypocotyl cells, while *p35S::CIP7-YFP*, *p35S::YFP-CIP7* and *pCIP7::CIP7-YFP* labeled cytoskeleton-like structures in hypocotyl epidermal cells (Fig. 3a).

**Fig. 3.**
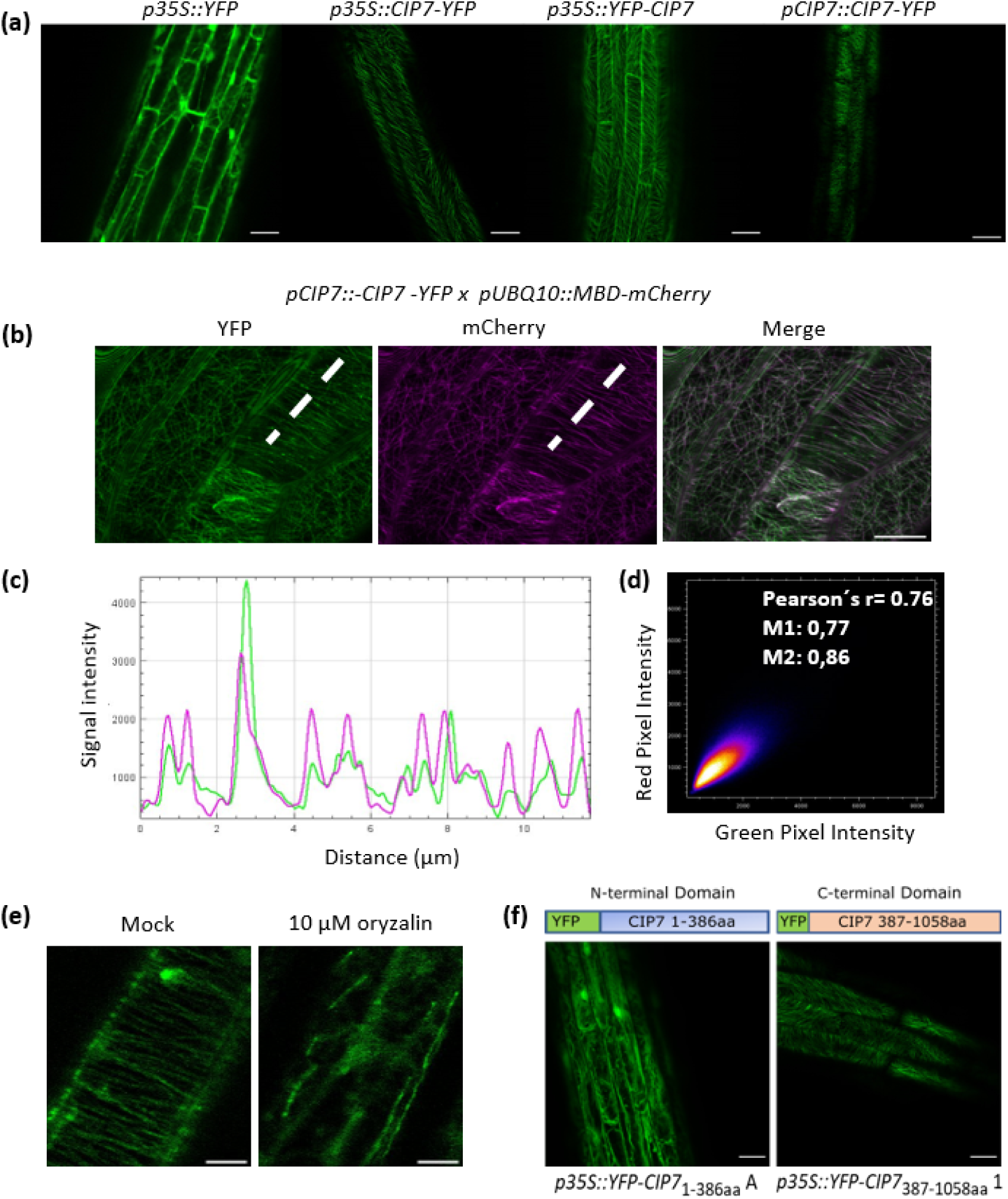
CIP7 localizes to cortical MTs in hypocotyl cells. **(a)** CIP7 localizes to cytoskeleton’s structures in the dark. Confocal microscopy images that show epidermal cells of hypocotyls from 5-d-old dark-grown seedlings. YFP fluorescence signal in green. *p35S::YFP*: Transgenic line expressing YFP driven by 35S promoter. *p35S::CIP7-YFP* and *p35S::YFP-CIP7:* Transgenic lines expressing the fusion proteins CIP7-YFP and YFP-CIP7 respectively, under the control of 35S promoter. *pCIP7::CIP7-YFP:* Translational reporter with native promoter. Scale bar: 25 μm. **(b)** Confocal images with ZEISS airyscan of *pCIP::CIP7-YFP* crossed with the MT marker *pUBQ10::MBD-mCherry* line showing epidermal cells of hypocotyls from 5-d-old dark-grown seedlings. In green, YFP signal. In magenta, mCherry signal. Merge in White. The ROI (Region of interest) is displayed with a dashed White line. Scale bar (full white line): 10 μm. (**c)** Fluorescence intensity profiles for green (YFP) and magenta (mCherry) channels along the ROI depicted in B. **(d)** Scatter plots of the intensity registered in the green and magenta channels in B. The YFP signal is shown on the x axis, whereas the y axis corresponds to the intensity of mCherry channel. Individual dots in the scatter plot correspond to single pixels of the original picture. The colour code highlights the frequency of dots present in a certain region of the scatter plot (from blue to yellow and white with increasing frequencies). Pearson’s r coefficient and Mandeŕs coefficients M1 and M2 of colocalization are indicated in the top right corner of the plot. **(e)** Effect of oryzalin-induced MT depolymerization on CIP7-YFP localization. 5-d-old dark-grown *p35S::CIP7-YFP* seedlings were incubated in 10 μM oryzalin for 30 minutes. Confocal images showing epidermal cells of hypocotyls. CIP7-YFP signal in green. Scale Bar: 5 μm. **(f)** The C-terminal domain of CIP7 is necessary for MTs localization. Localization of CIP7 domains, left the N-terminal domain of 386aa and right the C-terminal domain of 672aa in epidermal cells of hypocotyls. Only the C-terminal domain localize to cortical MTs. Confocal images of 3-d-old dark-grown transgenic lines carrying the constructs depicted in the diagrams (the diagrams are not to scale). Scale bar: 25 μm.

To study if CIP7 localizes specifically to MTs we performed co-localization experiments by crossing the *pCIP7::CIP7-YFP, p35S::CIP7-YFP*, *p35S::YFP-CIP7* and *p35S::YFP* lines with plants carrying the *pUB10::MBD-mCherry* MT marker (Ivanov & Harrison, 2014). YFP fluorescence associated with CIP7 fusion proteins overlapped with the mCherry signal from the MTs marker (Fig. 3b,c,d and Fig. S4a,b,c). No overlap was observed between *p35S::YFP* control plants and *pUB10::MBD-mCherry* lines (Fig. S4a). To further confirm that CIP7 localization was associated to MT; incubation of transgenic *p35S::CIP7-YFP* dark-grown seedlings with the MT-disrupting drug oryzalin, induced delocalization of the YFP fluorescence on MTs (Fig. 3e). Interestingly, 18 out of the 24 MTs enriched motifs were found in the C-terminal domain (Fig. S3a). To evaluate the contribution of each domain we generated transgenic plants overexpressing YFP fusions to the N-terminal domain (*p35S::YFP-CIP7_1-386aa_*) or C-terminal domain (*p35S::YFP-CIP7_387-1058aa_*). C-terminal domain fusions localized to MTs of 3-d-old dark-grown hypocotyls cells, while N-terminal domain fusions localized to cytosol and nuclei (Fig. 3f). This result was confirmed with a second independent line for each domain (Fig. S4d). All these results show that CIP7 localizes to cortical MTs and that the C-terminal domain is sufficient for proper CIP7 localization.

### Light reduces CIP7 expression

CIP7 was described as a transcription factor induced by light (Yamamoto *et al*., 1998), so we studied GUS activity in seedlings exposed to longer periods of light. Three day-old dark-grown seedlings showed high GUS activity, but this activity was markedly reduced after 8h of WL and was almost undetectable in seedlings grown for 3d under continuous WL (Fig. 4a). Staining patterns and responses to light treatment were confirmed on a second, independent transcriptional reporter line (Fig. S5). These results contrast with those reported by Yamamoto et al. To validate our results, we decided to quantify *CIP7* relative transcript levels by qRT-PCR. WT seedlings grown under continuous WL showed approximately an 80% reduction in *CIP7* transcript levels compared to that expressed in etiolated seedlings (Fig. 4b). Moreover, *CIP7* expression in translational reporter lines (*pCIP7::CIP7-YFP* in WT background) grown in the light showed a 50% reduction of total *CIP7* expression (Fig. 4b). CIP7 localization in 3-d-old light-grown *p35S::CIP7-YFP* and *p35S::YFP-CIP7* lines display fluorescence in cytoskeleton-like structures of hypocotyls and cotyledons including stomata (Fig. S6a). No localization within the nucleus was observed (Fig. S6b). YFP fluorescence levels in *pCIP7::CIP7-YFP* quantified by confocal microscopy showed significantly reduced levels in light-grown transgenic lines compared to dark-grown ones (Fig. 4c), according to GUS staining patterns and qRT-PCR results. Altogether, our results suggest that CIP7 is expressed in dark, and that its expression decays after at least 8h of exposure to WL.

**Fig. 4.**
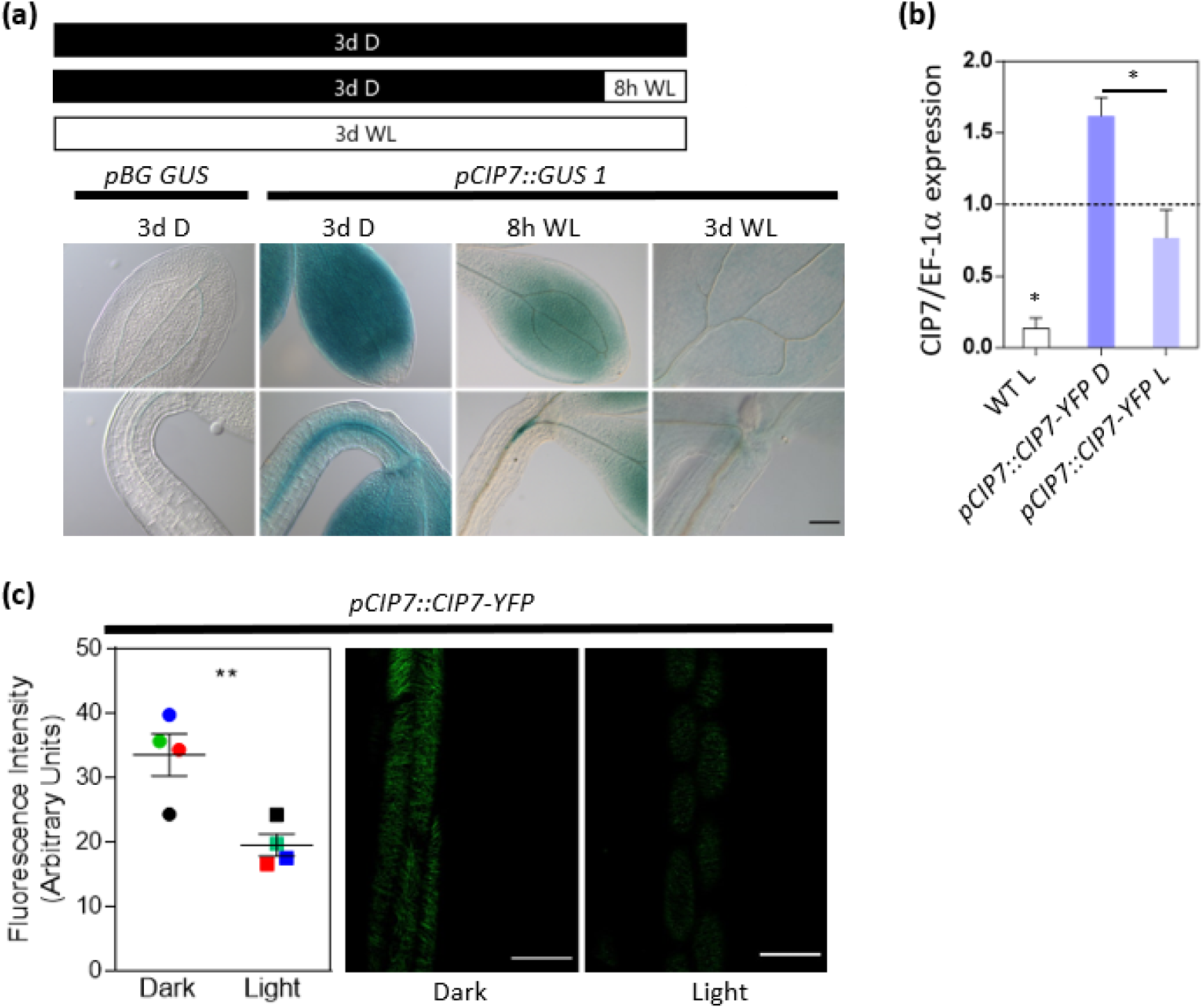
*CIP7* is expressed in darkness and decays with long white-light exposure **(a)** GUS expression pattern of 3-d-old *pCIP7::GUS* seedlings in different light treatments. Upper: Scheme of treatments: black box corresponds to dark, white box corresponds to white-light treatments; Down: GUS expression patterns of *pCIP7::GUS*. The assay was repeated twice with 2 different independent transgenic lines (see Fig. S5). Scale Bar: 100 μm. **(b)** Quantification of *CIP7* transcripts levels by RT-qPCR in WT and *pCIP7::CIP7-YFP*. Dashed line represents relative expression levels in WT 3-d-old dark-grown seedlings normalized to housekeeping gene EF-1alpha. WT L: 3-d-old light-grown WT seedlings. *pCIP7::CIP7-YFP* D and L: 3-d-old dark-grown (D) and light-grown (L) transgenic seedlings expressing fusion protein CIP7-YFP under control of the native promoter. The assay was performed twice. The plot depicted mean ±SE. Asterisks denote p-value < 0.05. **(c)** Quantification of fluorescence intensity of CIP7-YFP fusion protein in 3-d-old dark-grown and light-grown *pCIP7::CIP7-YFP* seedlings. Left, plot depicting mean ± SE of 4 experiments with 3-5 plants each. Three-six cells per plant and 3 measurements per cell were analyzed by ImageJ. Asterisks denote p-value < 0.01. Right, representative confocal images showing hypocotyls of 3-d-old dark or light-grown *pCIP7::CIP7-YFP* seedlings. Scale Bar: 40 μm.

### CIP7 affects hypocotyl elongation and the orientation of cortical MTs in dark-grown hypocotyl cells

To determine the biological function of CIP7, we generated *cip7* mutants by the CRISPR/Cas9 system. We obtained two independent lines *cip7_ed1* and *cip7_ed2* mutants. *cip7_ed1* presented a deletion of 89 amino acids in the N-terminal domain that produced a protein that maintained the frame. *cip7_ ed2* mutants presented an insertion of a T nucleotide producing a frameshift that introduces a premature stop codon, thus, generating a protein with 142 amino acids (Fig. 5a). MAPs mediate hypocotyl cell elongation (Sambade *et al*., 2012; Lian *et al*., 2017; Ma *et al*., 2018) and CIP7 is abundant in darkness (Fig. 4). After 3d in dark, *cip7_ed2* mutants showed reduced hypocotyl elongation compared to the WT, while *cip7_ed1* mutants were not different (Fig. 5b). This was not surprising as *cip7_ed1* mutants generate a protein with an intact C-terminal domain that shows MTs localization (Fig. 3f and Fig. S4d). 3-d-old light-grown *cip7_ed2* mutant seedlings were similar to the WT (Fig. S7).

**Fig. 5.**
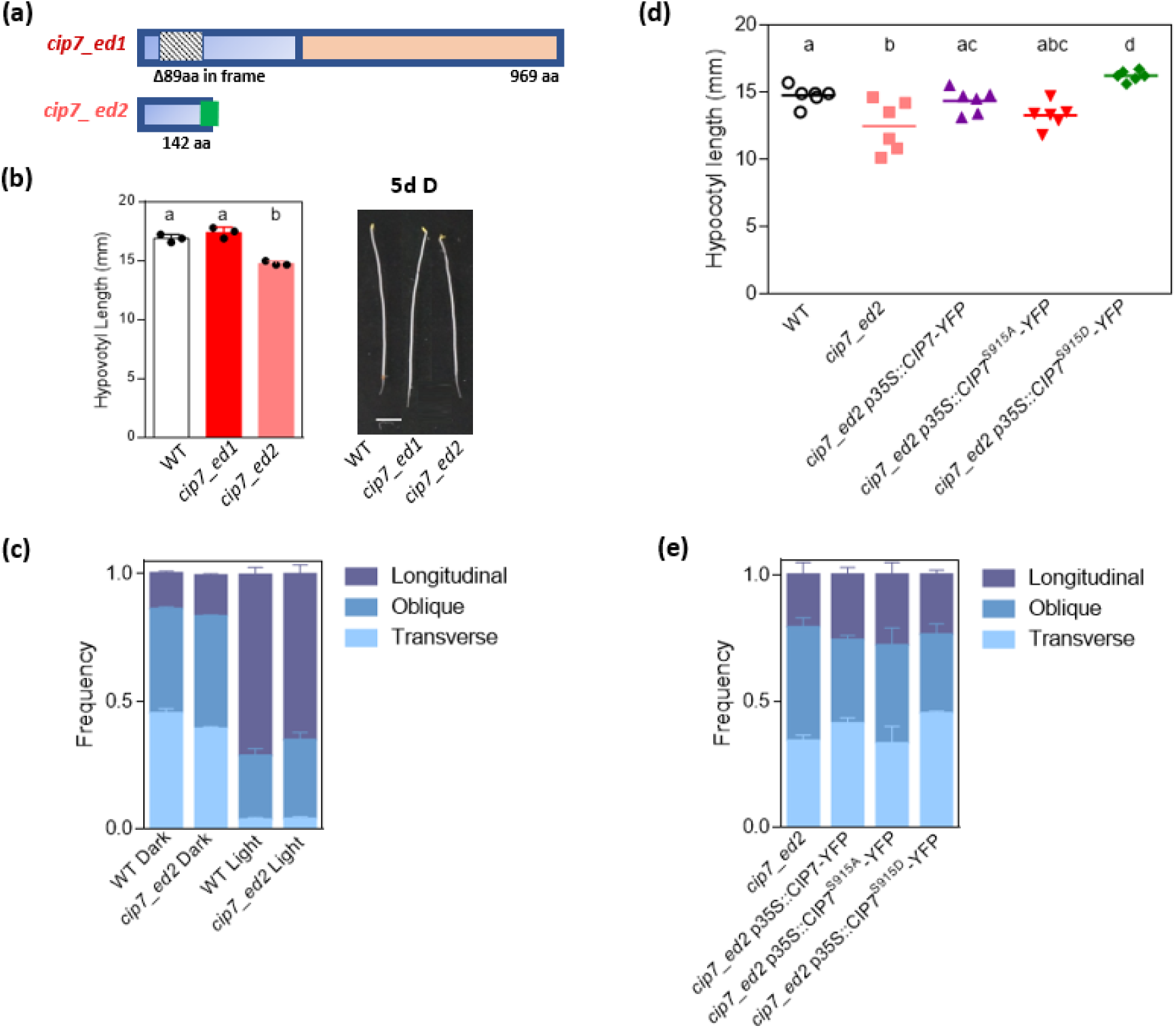
CIP7 effects on hypocotyl growth and the orientación of cortical MTs in the dark-grown hypocotyl cells are dependent on Sp^915^ isoform. **(a)** Scheme representing mutations in *CIP7* obtained by CRISPR/Cas9 edition. *cip7_ed1*: *cip7* mutant with a 89aa, in-frame deletion. *cip7_ed2*: knock out mutant with a premature stop codon. **(b)** Hypocotyl length of 3-d-old dark-grown seedlings of *cip7_ed1* and *cip7_ed2* mutants. At right, image of representative dark-grown seedlings. Scale bar: 5mm. The plot depicts media ±SE of 3 independent replicates for each genotype. Each replicate is the average of hypocotyl length measurements from 10 seedlings. One way ANOVA, Different letters show statistically significant differences. (p<0,05). **(c**) Frequency of epidermal hypocotyl cells from WT seedlings (*pUB10::MBD-mCherry*) and *cip7_ed2* mutants; that exhibit the MTs organization classes: Transverse, Obliques and Longitudinal. Seedlings were grown for 3 d in complete darkness or exposed to 20 min of WL. Data represents 11 independent experiments. Data was analyzed by ANOVA. *cip7_ed2* was backcrossed with the pUB10::MBD-Cherry line. **(d**) Hypocotyl length of 3-d-old dark-grown seedlings corresponding to WT, *cip7_ed2* and *cip7_ed2* complemented lines (*cip7_ed2 p35S::CIP7-YFP*, *cip7_ed2 p35S::CIP7^S915D^-YFP* carrying a phosphomimic isoform and *cip7_ed2 p35S::CIP7^S915A^-YFP* carrying a phosphonull isoform). The plot depicts measurements of hypocotyl length, mean± SE of 6 independent replicates for each genotype. Each replicate is the average of hypocotyl length measurements from 10 seedlings. Significance was evaluated by one way ANOVA. Different letters show statistically significant differences (p<0,05). **(e)** Frequency of epidermal hypocotyl cells from 3 d dark-grown seedlings of the *cip7_ed2* mutants and *cip7_ed2 p35S::CIP7-YFP*, *cip7_ed2 p35S::CIP7^S915A^S-YFP*, *cip7_ed2 p35S::CIP7^S915D^-YFP*, that exhibit the MTs organization classes: Transverse, Obliques and Longitudinal. Data represents the mean ±SE of 8 independent experiments. Data was analyzed by ANOVA. All mutants were backcrossed with the *pUB10::MBD-Cherry* line.

It has been reported that MT orientation is related to cell growth (Li *et al*., 2011; Lloyd, 2011; True & Shaw, 2020). Because our observations showed that CIP7 localizes to MTs and promotes hypocotyl elongation in the dark, we investigated MTs orientation in dark-grown seedlings. MT angles were categorized for each cell into transverse (0°-30°), oblique (30°-60°) and longitudinal (60°-90°) in relation to the long axis of the cell (Adamowski *et al*., 2019) (Fig. S8). Those cells that showed MTs in multiple orientations (random) were not considered (Adamowski *et al*., 2019). To analyze MT orientation, the MT marker was introduced into *cip7_ed2* mutant background by crossing. In 3-d-old dark-grown seedlings, the frequency of transverse MTs in *cip7_ed2* mutants was significantly lower than WT (p < 0,01), whereas the frequency of oblique and longitudinal oriented MTs was higher than WT controls (p < 0,05) (Fig. 5c). This distribution was expected for a genotype with shorter hypocotyl length. To test if CIP7 has a role in the dynamic reconfiguration of the cytoskeleton by light, we quantified MT orientation after 20 min of light. Exposure to light decreased transverse (p < 0.01) and increased longitudinal (p < 0.01) MTs orientation in the WT (Fig. 5c). MT orientation in *cip7_ed2* mutant hypocotyl cells was similar to that of the control after 20 min of WL, thus CIP7 does not mediate MTs rearrangements in light.

### CIP7 activity depends on Sp^915^ phosphorylation

To study if the phosphorylation in Sp^915^ is relevant for hypocotyl growth regulation we generated the phosphomimetic (substitute Serine by Aspartic acid, p*35S::CIP7^S915D^-YFP*), the phosphonull (change Serine to Alanine, p*35S::CIP7^S915A^-YFP*) and the control *p35S-CIP7-YFP* transgenic line versions in the *cip7_ed2* mutant background. We selected transgenic lines that express similar YFP levels among all (Fig. S9). Three-day-old dark-grown *cip7_ed2 p35S-CIP7-YFP* transgenic lines were able to rescue *cip7_ed2* mutant phenotype (Fig. S5d). Also, *cip7_ed2 p35S::CIP7^S915D^-YFP* seedlings were significantly taller than *cip7_ed2 p35S-CIP7-YFP* while *cip7_ed2 p35S::CIP7^S915A^-YFP* were not different from *cip7_ed2* mutants (Fig. S5d). Our data suggest that phosphorylated CIP7 in Sp^915^ promotes hypocotyl elongation in dark.

We then analyzed MT orientation in phosphomimetic and phospho-null dark-grown transgenic lines in the *cip7_ed2* mutant background. In darkness, *cip7_ed2 p35S::CIP7-YFP* showed a higher frequency of transverse MTs (p < 0.05) and lower frequency of oblique (p < 0.01) MTs compared to *cip_ed2* lines (Fig. 5e). In the same trend, *cip7_ed2 p35S::CIP7^S915D^-YFP* transgenic lines displayed a significant increase in transverse MTs (p < 0.05) and a decrease in oblique ones (p < 0.05) compared to *cip2_ed2 p35S::CIP7-YFP* (Fig. 5e). As expected from our previous results, *cip7_ed2 p35S::CIP7^S915A^-YFP* seedlings were not different from *cip7_ed2* mutants (Fig. 5e). This result is consistent with CIP7-dependent MT orientation relying on Sp^915^ phosphorylation. The non-phosphorylated isoform of CIP7 however, might not affect MTs binding capacity because *p35S::CIP7^S915A^-YFP* still localizes to MTs (Fig. S10).

## Discussion

In this study, we show that a small repertory of proteins undergoes early-light induced changes in their phosphorylation pattern and that MAPs are highly overrepresented in this group (Table 1). Furthermore, we show that among these MAPs, CIP7 promotes hypocotyl growth and transverse orientation of MTs in etiolated hypocotyl cells (Fig. 5b,c), a function that depends on Sp^915^ phosphorylation (Fig. 5d,e). CIP7 Sp^915^ isoform is rapidly reduced by light-activated sensory receptors (Fig 2a,b) and longer light exposures decrease *CIP7* expression (Fig. 4).

Phosphorylation of MAPs is a well-known key regulatory mechanism during neuronal activity that can affect MT affinity, cellular localization, or its overall function (Ramkumar *et al*., 2018; Barbier *et al*., 2019). The protein tau, a neuronal MAP in the central nervous system, undergoes multiple phosphorylation events and aggregation that are pathological hallmarks of Alzheimer disease and other tauopathies (Noble *et al*., 2013; Wegmann *et al*., 2021; Stefanoska *et al*., 2022). In plants, phosphorylation of the MAP SPR1 (SPIRAL 1) at Sp^6^ induces its disassociation from MTs favouring MTs stability (Wang *et al*., 2023). The protein phosphatase PP2A dephosphorylates microtubule-severing enzyme KATANIN to promote the formation of circumferential cortical MTs arrays in conical cells (Ren *et al*., 2022). Finally, phosphorylation at Sp^670^ of MICROTUBULE ASSOCIATED STRESS PROTEIN 1 (MASP1) is required for its function (Badiger Bhaskara *et al*., 2017).

Some MAPs promote while others inhibit hypocotyl cell elongation (Buschmann & Lloyd, 2008; Li *et al*., 2011; Wang *et al*., 2012). However, regulation of these MAPs by phosphorylation remained largely unexplored. MAPs regulation by phosphorylation does occur during plant cell division (Vavrdová *et al*., 2019; Motta & Schnittger, 2021), but post-germination hypocotyl growth depends on cell growth rather than cell division (Gendreau *et al*., 1997; Boron & Vissenberg, 2014; Daher *et al*., 2018). The reorganization of MTs in response to light occurs after 15 min (Lindeboom *et al*., 2019), 2019), thus the timing of light-induced changes in MAPs phosphorylation in our study is consistent with a regulatory role during early de-etiolation. The role of Sp^915^ isoform of CIP7 in the regulation of MT orientation and hypocotyl promotion supports this idea. Our work gives strong evidence that CIP7 localizes to MTs in the dark and when plants are transitioned to light. First *pCIP7::CIP7-YFP*, *p35S::CIP7-YFP*, *p35S::YFP-CIP7*, *p35S::CIP7^S915D^-YFP* and *p35S::CIP7^S915A^-YFP* transgenic lines localized to cytoskeletal structures (Fig. 3a and Fig. S6). Second, *pCIP7::CIP7-YFP*, *p35S::CIP7-YFP*, *p35S::YFP-CIP7* colocalized with MTs marker transgenic line (Fig 3b,c,d and Fig. S4a,b,c). Third, depolymerization of MTs with oryzalin disrupted *CIP7-YFP* localization (Fig. 3e). Fourth, 24 MTs binding motifs could be predicted using the bioinformatic tool *MAPanalyzer* (Fig. S3) and the C-Terminal domain of CIP7 containing 18 MTs binding motifs is necessary for MTs association (Fig. 3f and Fig. S4d). This result contrasts with that reported before, where transgenic lines carrying a chimeric genome/cDNA of CIP7 fused to GFP show GFP signals in the nucleus of dark-grown seedlings (Yamamoto *et al*., 1998). However, nuclear localization could not be completely discarded in other conditions. For example, the animal transcription factor MIZ-1 (MYC INTERACTING ZINC FINGER PROTEIN 1) binds to MTs in cytoplasm (Krtková *et al*., 2016) but when MTs are depolymerized MIZ-1 is released and imported to the nucleus to regulate gene expression (Ziegelbauer *et al*., 2001). CIP7 has been proposed to be a transcription factor, however CIP7 lacks a recognizable DNA binding domain (Yamamoto *et al*., 1998). Further studies are required to determine if CIP7 can act as a multifunctional MAP (Krtková *et al*., 2016).

Hypocotyl growth is inhibited in dark-grown knockout *cip7_ed2* mutant seedlings; and *cip7_ed2 p35S:CIP7-YFP* and phosphomimetic *cip7_ed2* p35S::CIP7^S915D^-YFP transgenic lines rescue the mutant phenotype (Fig. 5b,d) while phospho-null *cip7_ed2 p35S::CIP7^S915A^-YFP* line was similar to *cip7_ed2* mutant. Immediately after germination in dark, hypocotyls of Arabidopsis grow acropetally by the extension of its basal cells (Gendreau *et al*., 1997; Boron & Vissenberg, 2014). After 3-4 days post-germination basal cells stop growing and apical cells begin their expansion (Gendreau *et al*., 1997). GUS reporter-based analysis of the CIP7 promoter showed expression in apical regions of the hypocotyl (Fig. 2c and 4a) that is compatible with the pattern expression of proteins with a positive regulatory role in hypocotyl growth (Li *et al*., 2011).

The promotion of hypocotyl growth in darkness by CIP7 could be, at least in part, explained by the effects of the Sp^915^ isoform on MTs orientation. *cip7_ed2* mutants showed reduced hypocotyl length and a bias of oblique and longitudinal MT orientation pattern in detriment of transverse orientation (Fig. 5b,c). Accordingly, complementation of *cip7_ed2* mutants with *p35S-CIP7-YFP*, increases hypocotyl length and the frequency of transverse MTs (Fig. 5d,e), and complementation with cip*7_ed2 p35S::CIP7^S915D^-YFP* increased hypocotyl length and have more transverse MTs than *cip7_ ed2* mutants and also *cip7_ed2 p35S-CIP7-YFP* lines (Fig. 5d,e). The N-terminal domain that contains 6 from the 24 predicted MT binding motifs is not necessary for CIP7 MTs localization (Fig. 3f and Fig. S4d), and it also appears to be dispensable for CIP7 to fulfil its function as a growth promoter, because *cip7_ed1* mutants that retains a complete C-terminal domain but with a deletion in the N-terminal domain do not present hypocotyl phenotype in darkness (Fig. 5b).

Particularly, the phosphopeptide that contains Sp^915^ of CIP7 is present in dark-grown seedlings and *phyA phyB cry1 cry2* mutants after 20 min of white light, but absent in WT samples. CIP7 GUS pattern expression is similar in complete dark-grown seedlings with or without a 20 min light pulse (Fig. 2c). The reduced levels of Sp^915^ isoform of CIP7 after a short light pulse in the presence of phyA, phyB, cry1 and or/cry2 might be due to photoreceptor-mediated dephosphorylation (Yue et al., 2016 Kim et al., 2002 Moller et 2003). The detection of other phosphopeptides of CIP7 after the light pulse is not compatible with a photoreceptor-mediated degradation of the phosphorylated isoform (Hardtke *et al*., 2000; Duek *et al*., 2004; Jang *et al*., 2008; Sarid-Krebs *et al*., 2015; Wang *et al*., 2021). Specific CIP7 dephosphorylation after a short light signal might be a fast-acting mechanism that contributes to decrease hypocotyl growth.

In summary, this report gives evidence that phosphorylation of MAPs is relevant in the early steps of de-etiolation and our study on CIP7 and its phosphorylation adds new roles for this protein in the regulation of microtubule-based processes in the control of hypocotyl elongation. The role of cortical MTs in hypocotyl length during de-etiolation remains unclear and regulation by phosphorylation of MTs in the plant cell growth regulation needs to be further studied.

## Acknowledgements

We thank Dr. Jorge Casal (FAUBA-FIL, Argentina) for the critical reading of the manuscript and Dr. Pablo Pomata (IBYME, Argentina) for confocal microscopy support.

## Authors’ contributions

M.A.M, D.W., D.A and N.B. conceptualized the study. D.A, N.B, M.A.M and D.W. designed the experiments. M.A.M wrote the main manuscript text. D. A, N.B and D.W conducted the experiments. D.A; N.B; D. W and M.A.M analyzed data. All authors edited and approved the manuscript.

## Supplemental Material

**Fig. S1. Large scale phosphoproteome experimental design.**

**Fig. S2. MAPs are significantly overrepresented after a white-light pulse compared to darkness.**

**Fig. S3. CIP7 predicted MTs binding domains**

**Fig. S4. CIP7-YFP and YFP_CIP7 colocalize with cortical MTs.**

**Fig. S5.GUS expression pattern of pCIP7::GUS does not change after 20 minutes of WL treatments, but decays after long exposure to light.**

**Fig. S6. CIP7 localizes to cytoskeleton’s structures also in the light. Fig S7. Hypocotyl length of 3-d light-grown seedlings.**

**Fig. S8. MTs orientation.**

**Fig. S9. Different transgenic lines display similar expression levels. Fig. S10. Phosphorylation In S915 does not change CIP7 localization.**

**Table S1. Primers.**

**Table S2. Vectors and resistance.**

