## Supplemental material for "Radip light-induced phosphorylation changes in microtubule related proteins in arabidopsis"

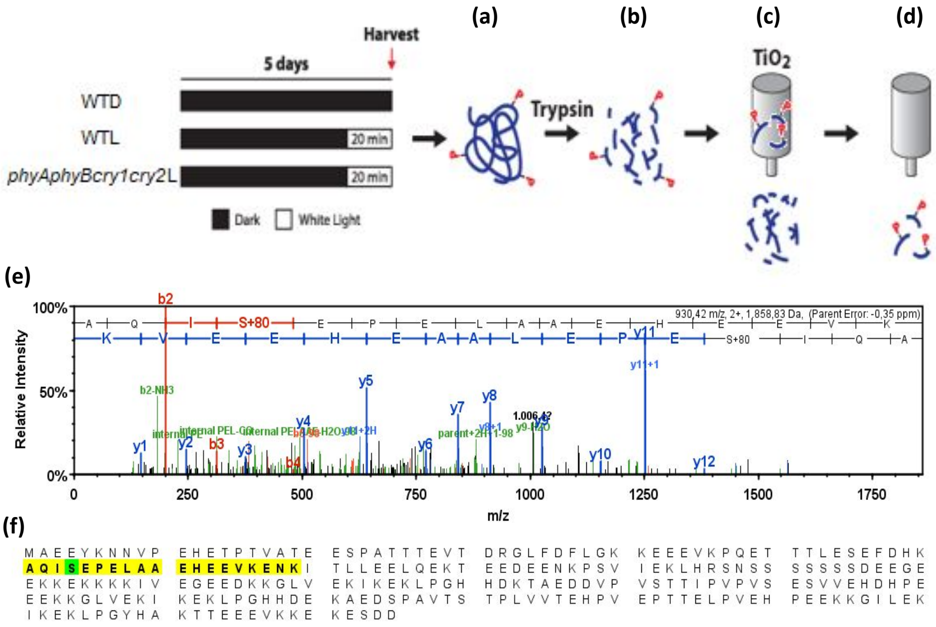


### Fig. S1.

**Large scale phosphoproteome experimental design.** Samples used for the phosphoproteomic analysis: WTD: 5-d-old dark-grown *Arabidopsis thaliana* WT seedlings; WTL: 5-d-old dark-grown WT seedlings exposed for 20 min to a WL pulse before harvest and *phyAphyBcry1cry2L:* 5-d-old dark-grown quadruple mutant seedlings exposed for 20 min to a WL pulse before harvest. Workflow: Total protein extraction were digested with trypsin to obtain peptides that were enriched in phosphopeptides with $TiO_2$ columns. After LC MS/MS we Identify phosphopeptides and phosphosites as is described in M&M. **(a)** Total protein extraction. **(b)** Trypsin digestion of proteins to obtain peptides. **(c)** Phosphopeptides enrichment. Phosphopeptides are retained in the column and non-phosphorylated peptides are discarded. **(d)** Elution of phosphopeptides. **(e)** LC-MS/MS. **(f)** Identification of phosphopeptides and phosphosites.

| | Total | Number with MAP motifs predicted | Number without MAP motifs predicted by *MAPanalyzer* | % with MAP motifs predicted | $X^2$/ p value |
|---|---|---|---|---|---|
| **A)** Genome | 27.417 | 1495 | 25922 | 5% | |
| **B)** Phosphoproteins identified in WTD | 933 | 130 | 903 | 14% | $X^2$= 125 p< $10^{-5}$ (A and B) |
| **C)** Phosphoproteins whose phosphorylation status differentially changed by light | 20 | 9 | 11 | 45% | $X^2$= 61; p<$10^{-5}$ (B and C) |

### Fig S2.

**MAPs are significantly overrepresented after a white-light pulse compared to darkness.** Frequency distribution of MAPs in the genome (A), the phosphoproteins identified in WTD (B) and the phosphoproteins that change significantly the phosphorylation pattern after 20 min of WL (C). Between brackets is represented the statistical comparison performed by $X^2$. A protein was considered MAP if it was reported with at least a 95% confidence by *MAPanalyzer*.

(a)

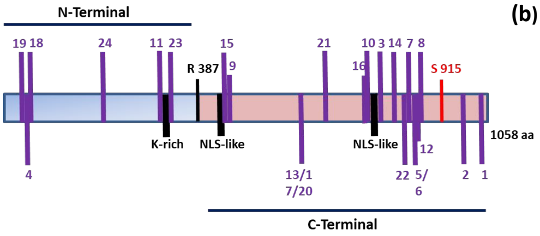


(b)

| Ref. | Motif | aa Start | aa End |
|---|---|---|---|
| 1 | H.R....S | 1029 | 1037 |
| 2 | TK..S | 973 | 978 |
| 3 | S.S.S...K | 788 | 797 |
| 4 | S...RPS | 59 | 66 |
| 5 | R..KP | 852 | 857 |
| 6 | RT..P | 852 | 857 |
| 7 | P..K.T | 848 | 854 |
| 8 | KPS..K | 855 | 861 |
| 9 | TS.....H | 437 | 445 |
| 10 | E..K.R.E | 763 | 771 |
| 11 | S.E...KK | 335 | 343 |
| 12 | SKPS | 854 | 858 |
| 13 | H..L...S | 601 | 609 |
| 14 | S..K..KT | 807 | 815 |
| 15 | HK.T | 434 | 438 |
| 16 | EEE.KK | 762 | 768 |
| 17 | H..L..S | 601 | 608 |
| 18 | RP.S...P | 63 | 71 |
| 19 | HL...S | 45 | 51 |
| 20 | HL...S | 603 | 609 |
| 21 | T..P..P | 676 | 683 |
| 22 | PK.....K | 841 | 848 |
| 23 | K.K.KQ | 350 | 356 |
| 24 | RLRE | 201 | 205 |

(c)

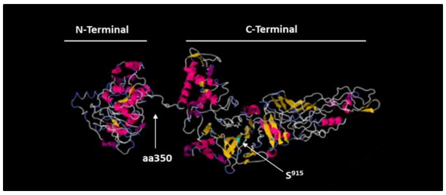


### Fig. S3.

**CIP7 predicted MTs binding domains. (a)** Diagram depicting the primary sequence of CIP7 protein and the location of the 24 putative MTs binding motifs consecutively numbered (in violet). CIP7 is a protein of 1058aa (amino acids) with 2 domains delimited by Arginine 387; containing K-rich (region rich in Lysines) and putative NLS-like motifs (nuclear localization signal) according to Yamamoto *et.al.*, 1998 description. The light-regulated phosphosite $S^{915}$ (in red) is located in the C-terminal domain. **(b)** Table specifying the sequence and accurate location of the 24 putative MTs binding motifs of CIP7 predicted with MAPanalyzer with a specificity higher than 95% (Zhou et al., 2015). **(c)** Model of CIP7 tertiary structure predicted by I-TASSER (Yang et al., 2015; Roy et al., 2010; Zhang, 2008). It shows the N-terminal domain of 350aa and the C-terminal domain of 672aa connected by a K-rich region.

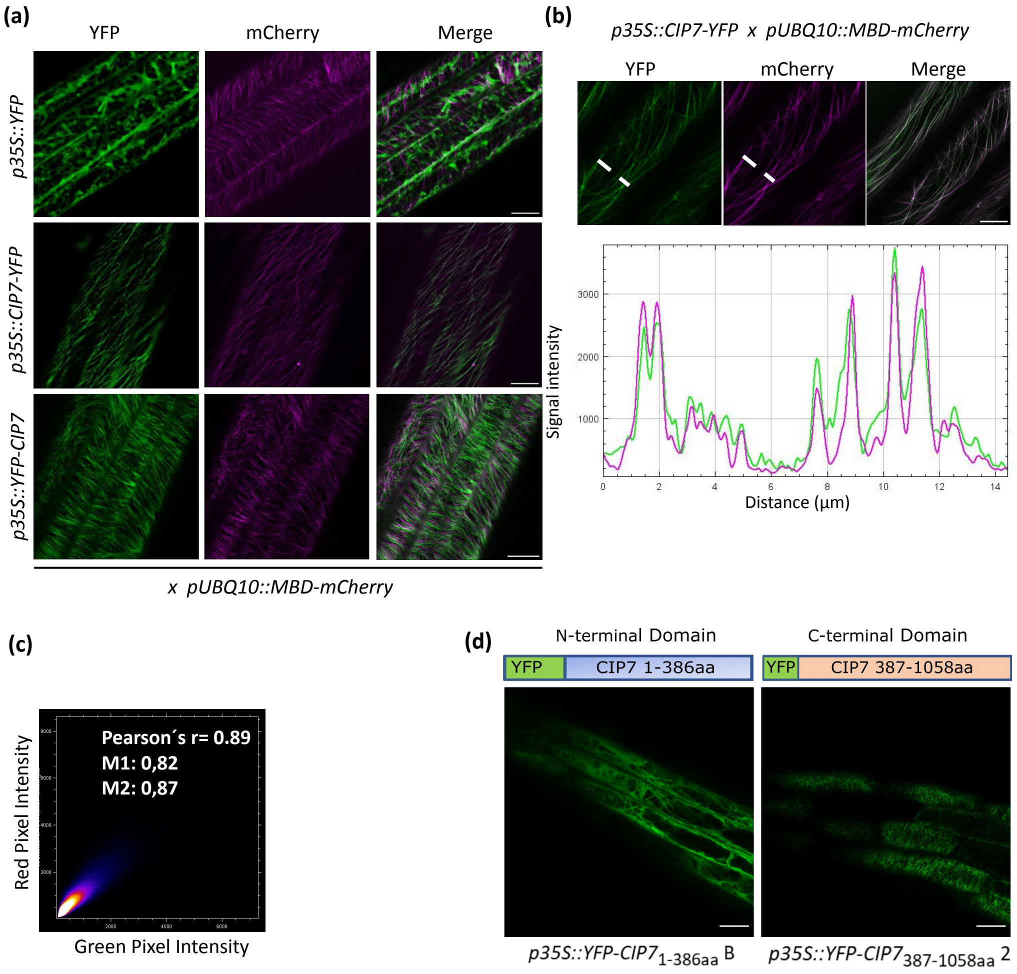


### Fig. S4.

**CIP7-YFP and YFP_CIP7 colocalizes with cortical MTs. (a)** Confocal images showing epidermal cells of hypocotyls from 5-d-old dark-grown seedlings. All the translational reporters of CIP7 (*p35S::CIP7-YFP* and *p35S::YFP-CIP7*) were crossed with MT marker transgenic line *pUBQ10::MBD-mCherry*. *p35S::YFP* was used as a control. In green, YFP signal. In magenta, mCherry signal. Merge in White. The experiment was repeated 3 times. Scale bar: 16 µm. **(b)** Upper: Confocal images with ZEISS airyscan of *p35S::CIP7-YFP* crossed with *pUBQ10::MBD-mCherry* line showing epidermal cells of hypocotyls from 5-d-old dark-grown seedlings. In green, YFP signal. In magenta, mCherry signal. Merge in White. The ROI (Region of interest) is displayed with a dashed White line. Scale: 10 µm. Down. Fluorescence intensity profiles for green (YFP) and magenta (mCherry) channels along the ROI. **(c)** Scatter plots of the intensity registered in the green and magenta channels in B. The YFP signal is shown on the x axis, whereas the y axis corresponds to the intensity of mCherry channel. Individual dots in the scatter plot correspond to single pixels of the original picture. The color code highlights the frequency of dots present in a certain region of the scatter plot (from blue to yellow and white with increasing frequencies). Pearson's r coefficient and Mander´s coefficients M1 and M2 of colocalization are indicated in the top right corner of the plot. **(d)** The C-terminal domain of CIP7 is necessary for MTs localization. Localization of CIP7 domains in a second independent transgenic lines. Only the C-terminal domain localize to cortical MTs. Confocal images of 3-d-old dark-grown transgenic lines carrying the constructs depicted in the diagrams. Scale bar: 25 µm

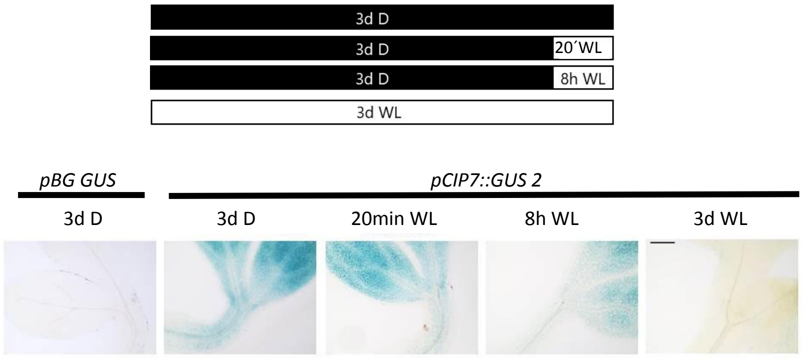


### Fig. S5.

**GUS expression pattern of *pCIP7::GUS* does not change after 20 minutes of WL treatments, but decays after long exposure to light**. GUS expression pattern of *pCIP7::GUS* 2 (independent line from that shown in Fig. 2c) compared to the no promoter control (pBG GUS) *in* 3-d-old dark-grown seedlings exposed or not to 20 min WL or 8 h of WL before harvest and in 3-d-old light-grown seedlings. Upper**:** Scheme of treatments: black box corresponds to dark, white box corresponds to light treatments; Down: GUS expression patterns of *pCIP7::GUS 2*.Scale Bar: 100 µm. pBG GUS corresponds to the non-promoter control.

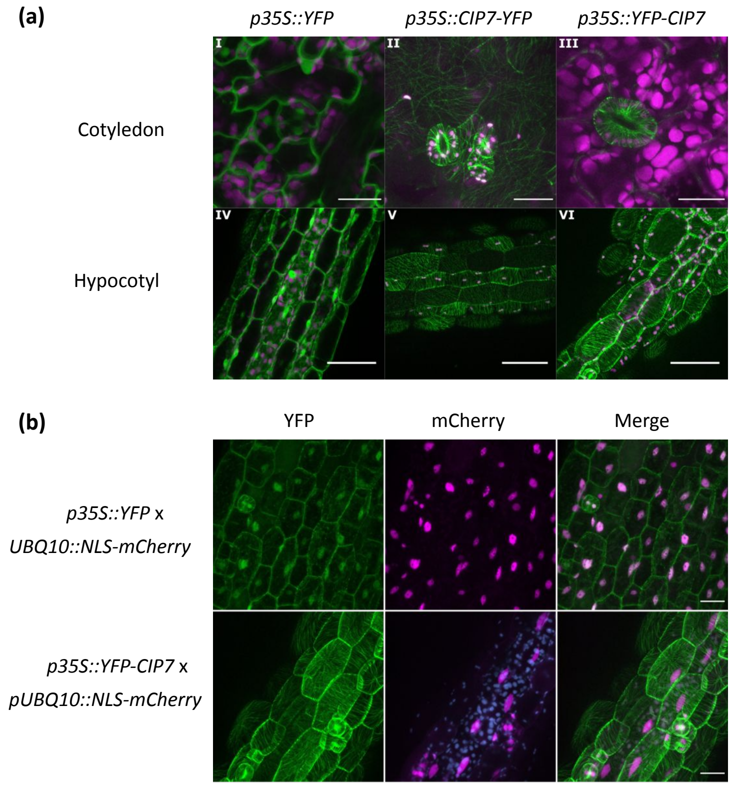


### Fig. S6.

**CIP7 localize to cytoskeleton's structures also in the light. (a)**. I- III Confocal microscopy images showing pavement cells of cotyledons from 3d light-grown seedlings. IV – VI Confocal microscopy images showing epidermal cells of hypocotyl from 3d light-grown seedlings. YFP fluorescence signal in green and chloroplasts in magenta. *p35S::YFP*: Transgenic line expressing YFP driven by 35S promoter. *p35S::CIP7-YFP* and *p35S::YFP-CIP7:* Transgenic lines expressing the fusion proteins CIP7-YFP and YFP-CIP7 respectively, under control of 35S promoter**. *pCIP7::CIP7-YFP:*** Transgenic lines expressing the fusion proteins CIP7-YFP under its native promoter. Scale bar in I, II, IV, V & VI: 50 µm. Scale bar in III: 30 µm. **(c)** Colocalization assays with the nuclear marker *UBQ10::NLS-mCherry* reveal CIP7 was not detected in nuclei. Confocal microscopy images showing epidermal cells of hypocotyl from 3d light-grown seedlings co-expressing YFP-CIP7 (green) and NLS-mCherry (magenta), showing no overlapping of YFP and mCherry signals. Seedlings *p35S:YFP* x *UBQ10::NLS-mCherry* were used as controls, showing white nuclei due to the overlapping YFP and mCherry signals. Chloroplasts in cyan. Scale bar: 25 µm.

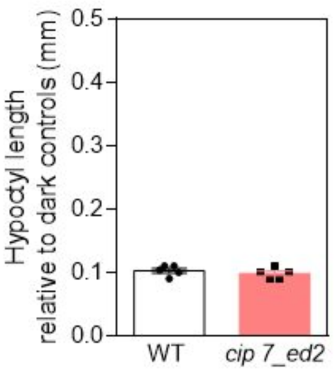


**Fig. S7.**

**Hypocotyl length of 3-d old light -grown seedlings.** Hypocotyl length relative to dark controls in WT and c*ip7_ed2* mutant seedlings grown for 3d in continuous light.

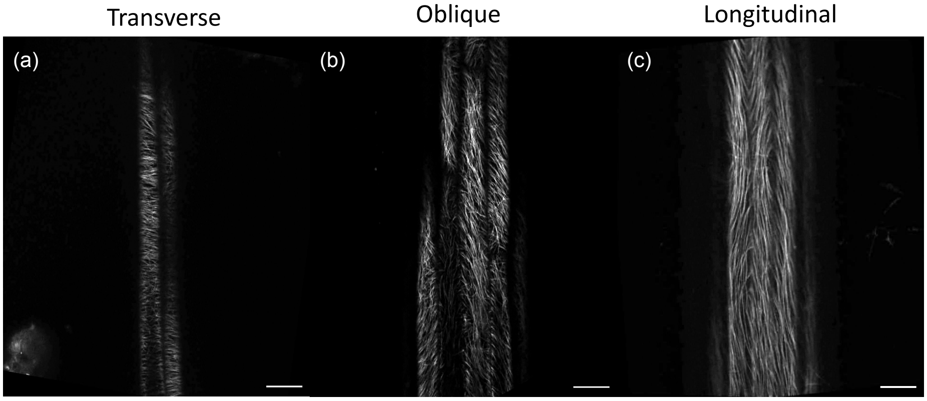


**Fig. S8.**

**MTs orientation.** Confocal microscopy images of 3-d-old dark-grown epidermal hypocotyl cells of MBD-mCherry expressing plants showing showing different MT orientation categories: transverse (Angle= 13°) **(a)**, oblique (Angle= 46°) **(b)** and longitudinal (Angle= 89°) **(c)** Scale bar: 25 µm.

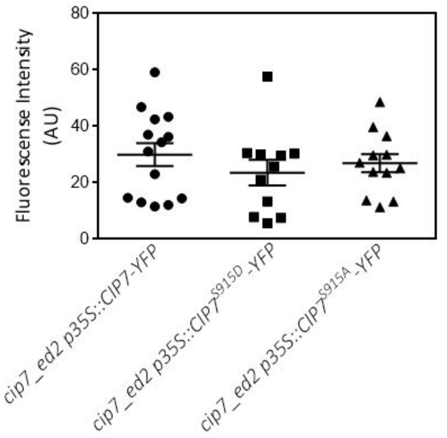


### Fig. S9.

**Different transgenic lines display similar expression levels.** Quantification of fluorescence intensity in 3-d-old dark-grown seedlings of the transgenic lines used in Figure 6C.

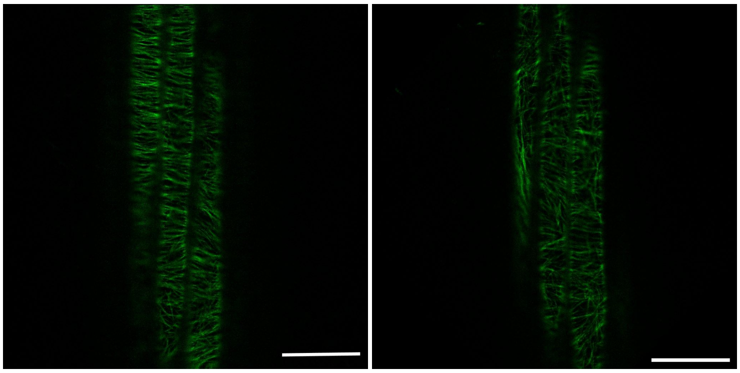

*cip7_ed2 p35s::CIP7$^{S915D}$-YFP* *cip7_ed2 p35s::CIP7$^{S915A}$-YFP*

### Fig. S10.

**Phosphorylation In $S^{915}$ does not changes CIP7 localization.** Confocal microscopy images of 3-d-old dark-grown epidermal hypocotyl cells in *cip7_ed2 p35s::CIP7$^{S915D}$-YFP* and *cip7_ed2 p35s::CIP7S$^{915A}$-YFP* lines. Scale bar right: 25 µm.

| Primer | Purpose | Sequence 5' - 3' |
|---|---|---|
| gCIP7F | Amplif. with NotI site | ATAT GCGGCCGC GCGAATCCTCTGGATGAGAC |
| CIP7pR | Amplif. with AscI site | ATAT GGCGCGCC TAAAGAACCCACAACAAGCAAAAAAA |
| cCIP7F | Amplif. with NotI site | ATAT GCGGCCGC A ATGGATCCAAGAACAAGACTTGAC |
| cCIP7-stop R | Amplif. with AscI site | ATAT GGCGCGCC C CTTTTTCCTTCCGAACATAAGAAGC |
| cCIP7+stop R | Amplif. with AscI site | ATAT GGCGCGCC TTACTTTTTCCTTCCGAACATAAGA |
| CIP7 1-386 R | Amplif. with AscI site | ATAT GGCGCGCC TTAGATAACCACTTTCCTGGATT |
| CIP7 387-1058 F | Amplif. with NotI site | ATAT GCGGCCGC A ATG CGCAACATAAATTATATAA |
| CIP7 300pb | Sequencing | GTGTGTCGGCGTACATAACG |
| CIP7 seq1 | Sequencing | GGAGATTGAGCAGATCGAGG |
| CIP7 seq2 | Sequencing | GGAAACTCTATGGATGCCTCG |
| CIP7 seq3 | Sequencing | GTGAGAAAGAGAGAACCGCC |
| YFP N seq | Sequencing | GCTGACCCTGAAGTTCATCTGC |
| YFP-Fw_seq | Sequencing | CGAGGTGAAGTTCGAGGGCGAC |
| CRISPR CR6 rv | Sequencing | GATGGAGACACTGTGGAAGA |
| CIP7 CR6 F | CAS9 guide | GATT **G**TATGTAGGAAACGAGTTGGG |
| CIP7 CR6 R | CAS9 guide | AAAC CCCAACTCGTTTCCTACATA**C** |
| RT-qPCR2 cip7 F | RT-qPCR | TCACCAGCGGTCTCAGAGAT |
| RT-qPCR2 cip7 R | RT-qPCR | CCAAGGCAACGGCTAAAGTC |
| EF-1$\alpha$_F | RT-qPCR | TGAGCACGCTCTTCTTGCTTTCA |
| EF-1$\alpha$_R | RT-qPCR | TGTAACAAGATGGATGCCACCACC |
| phosphoA_F | Substitution S > A | CTTCCTGCTGTTGCTGCTGTTGATGATTTCAAAG |
| phosphoA_R | Substitution S > A | CTTTGAAATCATCAACAGCAGCAACAGCAGGAAG |
| phosphoD_F | Substitution S > D | GTTGCTGATGTTGATGATTTCAAAGACA |
| phosphoD_R | Substitution S > D | ATCAACATCAGCAACAGCAGGAAGTGCATC |

| Amplicon | Combination of primers |
|---|---|
| Locus | gCIP7F + cCIP7-stop R |
| Promoter of 2500pb | gCIP7F + CIP7pR |
| Gene with stop codon | cCIP7F + cCIP7+stop R |
| Gene without stop codon | cCIP7F + cCIP7-stop R |
| Truncated version 1-386aa | cCIP7F + CIP7 1-386 R |
| Truncated version 387-1058aa | CIP7 387-1058 F + cCIP7+stop R |

### Table S1.

List of primers and the combinations used to obtain the different amplicons. The primers' names, purposes and sequences are specified.

| Vector | Construct | Resistence in *E. coli* (TOP10) | Resistence in *A. tumefasciens* (GV3101) | Resistence in Plant |
|---|---|---|---|---|
| pBG GUS | *pCIP7::GUS* | Spectinomicine 100mg/l | Rif. 50mg/l + Gent. 30mg/l + Spect. 100mg/l | BASTA 120µg/ml |
| pHGY | *pCIP7::CIP7-YFP* | Spectinomicine 100mg/l | Rif. 50mg/l + Gent. 30mg/l + Spect. 100mg/l | Hygromicine 50mg/l |
| pH35GY | *p35S::CIP7-YFP* | Spectinomicine 100mg/l | Rif. 50mg/l + Gent. 30mg/l + Spect. 100mg/l | Hygromicine 50mg/l |
| pH35YG | *p35S::YFP-CIP7* (& domains) | Spectinomicine 100mg/l | Rif. 50mg/l + Gent. 30mg/l + Spect. 100mg/l | Hygromicine 50mg/l |
| pH35GS | *p35S::YFP* | Spectinomicine 100mg/l | Rif. 50mg/l + Gent. 30mg/l + Spect. 100mg/l | Hygromicine 50mg/l |

**Table S2.**

List of vectors used for the different constructs. The antibiotics and concentrations used for selection in bacteria and plant are specified.
